# Joint contribution of adaptation and neuronal population recruitment to response level in visual area MT: a computational model

**DOI:** 10.1101/2024.03.13.584758

**Authors:** Maria Inês Cravo, Rui Bernardes, Miguel Castelo-Branco

**Affiliations:** Coimbra Institute for Biomedical Imaging and Translational Research (CIBIT), University of Coimbra, Coimbra, Portugal; Institute of Nuclear Sciences Applied to Health (ICNAS), University of Coimbra, Coimbra, Portugal; Faculty of Medicine, University of Coimbra, Coimbra, Portugal

## Abstract

Adaptation is a form of short-term plasticity triggered by prolonged exposure to a stimulus, often resulting in altered perceptual sensitivity to stimulus features through a reduction in neuronal firing rates. Experimental studies have explored adaptation to bistable stimuli, specifically a stimulus comprising inward-moving plaids that can be perceived as either a grating moving coherently downward or two plaids moving incoherently through each other. Functional magnetic resonance imaging (fMRI) recordings have shown higher activity during incoherent perception and lower activity during coherent stimulus perception. There are two potential explanations for the underlying neural mechanisms: a weaker coherent stimulus response may result from stronger adaptation to coherent versus incoherent motion, or a stronger incoherent stimulus response could stem from the involvement of more neural populations to represent motion in more directions. Here, we employ a computational model of visual neurons with and without firing rate adaptation to test these hypotheses. By simulating the mean activity of a network of thirty-two columnar populations of visual area MT, each tuned to one direction of motion, we investigate the impact of firing rate adaptation on the blood-oxygen-level-dependent (BOLD) signal generated by the network in response to coherent and incoherent stimuli. Our results replicate the experimental curves both during and after stimulus presentation only when the model includes adaptation, highlighting the importance of this mechanism. However, our findings reveal that the response to incoherent motion is larger than the response to coherent motion for a wide variety of stimulus parameters and adaptation regimes, suggesting that the observed reduced response to coherent stimuli is most likely due to the activation of smaller neuronal populations, in alignment with the second hypothesis. Hence, adaptation and differential neuronal recruitment work together to give rise to the observed hemodynamic responses. This computational work sheds light on experimental results and enriches our understanding of the mechanisms involved in neural adaptation, particularly in the context of heterogeneous neuronal populations.

## Introduction

Firing rate adaptation is a well-established feature of sensory neurons, involving short-term reduction in spiking activity and changes in sensitivity to the features of a repeatedly presented stimulus [1,2]. This process is an important resource to achieve code efficiency in the brain as it ensures heightened sensitivity to unexpected or unusual stimuli [3,4]. However, adaptation also leads to visual illusions such as the motion after-effect (MAE): the perception of motion in a stationary image following prolonged exposure to a moving stimulus [5]. This phenomenon is also known as the waterfall effect, coined after its notable observation in a natural setting.

In neuroscience research, visual adaptation has been documented through psychophysical experiments [6,7] and electrophysiological recordings [8,9]. In functional imaging, adaptation serves as a signature of region-specific encoding, with regions responsive to a specific feature displaying reduced activity [10–13]. Conversely, the search for a neural correlate of the MAE focused on increased activity in non-adapted neuron populations [14,15]. However, isolating the effects of adaptation is challenging, since functional magnetic resonance imaging (fMRI) shows the activity of entire voxels, likely containing diverse neuron populations coding for different features.

Adaptation is believed to play an important role in the perception of ambiguous stimuli by driving switches between the various available interpretations [16,17]. A notable bistable stimulus is the plaid stimulus, consisting of inward-moving plaids that can be perceived either as a grating moving coherently downward or as two plaids moving incoherently through each other. In an fMRI study, Sousa et al. [18] manipulated plaid stimuli to induce either coherent or incoherent perception, observing distinct levels of brain activation in the middle temporal (MT) visual area for each type of perception. Specifically, they noted that the response to coherent motion was weaker compared to the response to incoherent motion. There are two potential explanations for the underlying neural mechanisms (Fig. 1): a weaker coherent stimulus response may result from stronger adaptation to coherent versus incoherent motion, or a stronger incoherent stimulus response could stem from the involvement of more neural populations to represent motion in more directions.

**Figure 1.**
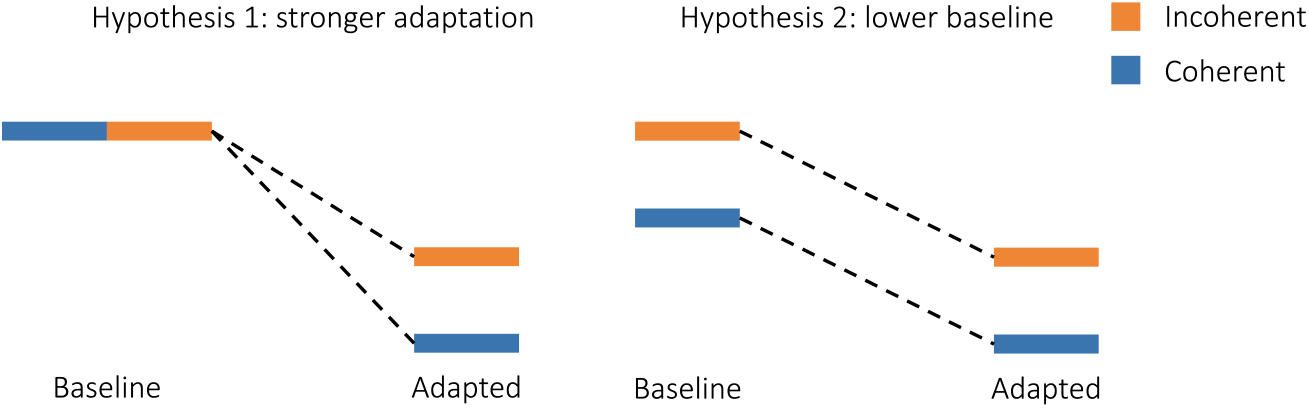
Alternative hypotheses to explain lower response to coherent motion. The first hypothesis proposes that incoherent and coherent stimuli elicit the same response, but the coherent representation undergoes a more pronounced reduction during prolonged exposure through adaptation. The second hypothesis proposes that the coherent stimulus elicits a smaller response due to recruitment of a smaller neural representation.

Here, we use a firing rate model and the Balloon-Windkessel model to simulate the activity and resulting blood-oxygen-level-dependent (BOLD) signal of a network of thirty-two neurons, each tuned to one direction of motion. By varying the levels of adaptation, we determine that strong adaptation is necessary to replicate the experimental curves, both during and after stimulus presentation. However, increasing adaptation does not affect the relative order of the levels of the coherent and incoherent response curves, suggesting that differences in neuronal recruitment are responsible for the lower response to coherent motion.

## Methods

To simulate the activity of area MT, we used a firing rate model with 32 neuronal units. Each unit represents a neuron or a population of identical neurons tuned to motion in one direction. The dynamics of each neuron is described by two differential equations: one for the evolution of the firing rate as a function of the synaptic current and one for the accumulation of adaptation as a function of the firing rate. Firing rate activity is then transformed into a BOLD signal using the Balloon-Windkessel model [19].

### Receptive Field

Each neuron responds maximally to a preferred direction of motion according to the following receptive field [6,20]:

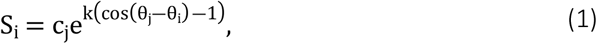

where S_i_ is the response of neuron i, with a preferred direction θ_i_, to a stimulus moving with intensity c_j_ in the direction θ_j_, and k is a measure of the bandwidth of the receptive field, with higher k corresponding to narrower tuning. For 32 neurons with equidistant preferred directions, θ_i_ = 0°,11.25°,12.5°,23.75°,35°,….

### Differential Equations

The synaptic current I entering each neuron is:

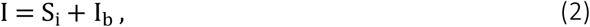

with S_i_ the sensory input, given by Eq. (1), and I_b_ the baseline activity. The synaptic process is assumed to be quasi-instantaneous compared to the firing rate process [21], which is described by:

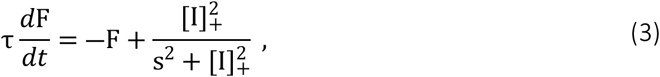

where F is the firing rate, τis the time constant of the low-pass filtering process, s is the saturation constant of the non-linear activation function and […]_+_ denotes half-wave rectification.

Firing rate adaptation is modeled as an exponential process [2,22]:

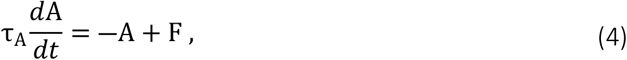

where *A* is the adaptation level of a given neuron, which accumulates with the firing rate *F* and decays with time constant *τ*_*A*_. Adaptation acts divisively on the activation function of each neuron, thereby modifying Eq. (3) to:

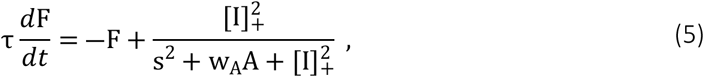

with *w*_*A*_ the adaptation strength. The activity of each neuron is thus calculated by integrating two differential equations, Eqs. (4) and (5).

### BOLD Signal

Neuronal activity gives rise to the blood-oxygen-level-dependent signal detected in fMRI through a hemodynamic process that is well-described by the Balloon-Windkessel model [19], comprised of four dynamical variables: vasodilatory signal s(t), blood inflow f(t), blood volume v(t) and deoxyhemoglobin content q(t). The system of differential equations is:

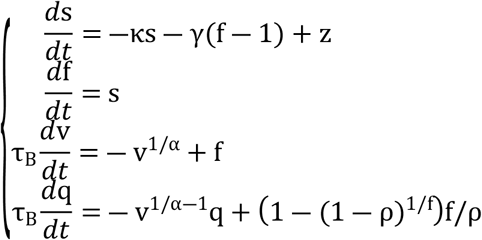

where z(t) is the neuronal activity, given by the sum of the neuronal firing rates calculated through Eq. (5), and κ, γ, τ _B_, α and ρ are parameters describing the rate of signal decay, rate of flow-dependent elimination, hemodynamic transit time, Grubb’s exponent of blood outflow and resting oxygen extraction fraction, respectively. The BOLD signal, B(t), is then a volume-weighted sum of intra- and extravascular contributions to blood volume and deoxyhemoglobin content:

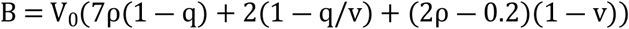

with *V*_0_ the resting blood volume fraction.

### Stimuli and Experimental Parameters

The bistable plaid stimulus is composed of two inward-moving plaids with two possible interpretations: a grating moving downward (coherent percept) or two plaids moving through each other (incoherent percept) (Fig. 2). In their study [18], Sousa et al. added a field of moving dots to the bistable plaid stimulus to induce the disambiguated perception of either coherent or incoherent motion. A non-adapting control stimulus was used, consisting of alternating coherent and incoherent motion in eight different directions. After 6 seconds of a static grating display, the stimulus was presented for 30 seconds, followed by 12 seconds of a static grating display. The plaids make a 60° static angle with the horizontal and move in the horizontal direction (left plaid moves to the right, 0°, and right plaid moves to the left, 180°). Besides the oriented lines that comprise each plaid, the background rhombi between the lines constitute an extra component of the stimulus. As such, the stimuli have motion energy in a maximum of three directions. The coherent stimulus is dominated by downward motion, while the incoherent stimulus includes motion in all three directions (see Supplementary Videos). Assuming the same total intensity for both coherent and incoherent conditions, the coherent stimulus may be defined as having intensity 1 in the downward direction (270°) and 0 in all other directions, while the incoherent stimulus can be defined as having intensity 1/3 in the 0°, 270° and 180° directions, and 0 in all other directions. These can be written as *s*_*c*_ = (0,1,0) for the coherent stimulus and *s*_*i*_ = (1/3, 1/3, 1/3) for the incoherent stimulus. The parametrization of the stimulus constitutes one of the main aspects of the model. These parameter values are summarized in Table 1.

**Table 1.** Stimulus parameters.

| Parameter | Value | Reference |
| --- | --- | --- |
| Direction of grating movement (°) | 270 | [18] |
| Direction of plaid 1 movement (°) | 0 | Ibid. |
| Direction of plaid 2 movement (°) | 180 | Ibid. |
| Onset of adapting stimulus (ms) | 6000 | Ibid. |
| Onset of testing stimulus (ms) | 36000 | Ibid. |
| Alternation duration in non-adapting condition (ms) | 1500 | Ibid. |
| Coherent stimulus intensity in grating direction | 1 | - |
| Coherent stimulus intensity in plaid directions | 0 | - |
| Incoherent stimulus intensity in grating direction | 1/3 | - |
| Incoherent stimulus intensity in plaid directions | 1/3 | - |

**Figure 2.**
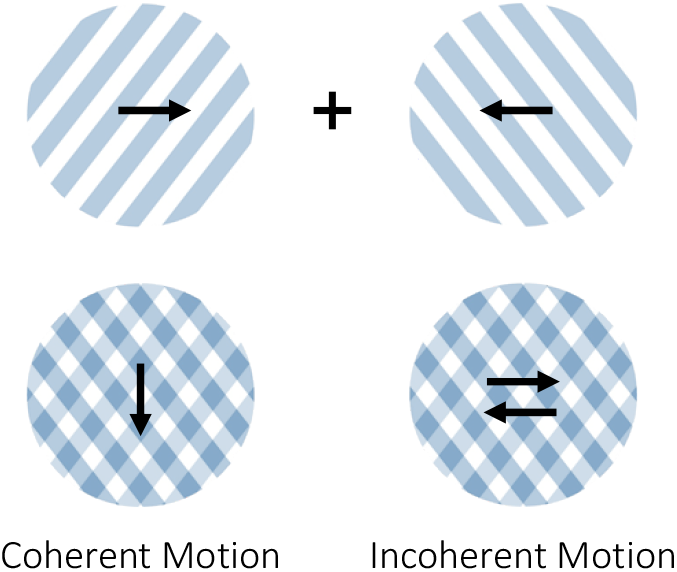
Bistable plaid stimulus: two superimposed plaids moving in opposite directions can be perceived as either one grating moving downward (coherent motion) or as two plaids moving past each other (incoherent motion).

### Simulation Details

The neuronal circuit was defined in PyRates [23,24], where the differential equations were solved via SciPy’s Runge-Kutta(4,5) integration scheme [25]. Parameter values are presented in Tables 2-4.

**Table 2.** Neuronal circuit parameters.

| Parameter | Description | Value | Reference |
| --- | --- | --- | --- |
| $k$ | Bandwidth of neuronal receptive field | 180 | - |
| $I_b$ | Baseline synaptic current | 0.1 | - |
| $\tau$ | Firing rate time constant (ms) | 50 | [22,26] |
| $s$ | Saturation of neuronal activation function | 0.5 | [22,26] |
| $\tau_A$ | Adaptation time constant (ms) | 2000 | [22,27,28] |
| $w_A$ | Adaptation strength | 0, 2, 4 | [22,27] |

**Table 3.** Hemodynamic parameters.

| Parameter | Description | Value | Reference |
| --- | --- | --- | --- |
| $\kappa$ | Rate of vasodilatory signal decay ($s^{-1}$) | 0.65 | [19,29,30] |
| $\gamma$ | Rate of flow-dependent elimination ($s^{-1}$) | 0.41 | Ibid. |
| $\tau_B$ | Hemodynamic transit time (s) | 0.98 | Ibid. |
| $\alpha$ | Grubb's exponent of blood outflow | 0.32 | Ibid. |
| $\rho$ | Resting oxygen extraction fraction | 0.34 | Ibid. |
| $V_0$ | Resting blood volume fraction | 0.02 | Ibid. |

**Table 4.** Simulation parameters.

| Parameter | Description | Value |
| --- | --- | --- |
| $T$ | Total integration time (ms) | 48000 |
| $dt$ | Integration time step (ms) | 0.1 |
| $dts$ | Sampling time step (ms) | 50 |
| $T_{init}$ | Initialization period, discarded (ms) | 6000 |

## Results

To elucidate the neuronal mechanisms probed by the fMRI experiment of Sousa et al. [18], we simulate the activity of thirty-two neuron populations, each with a preferred motion direction and an independent adaptation process. The total activity of the system is obtained by summing the firing rate of all units and the corresponding BOLD signal is calculated through the Balloon-Windkessel equations. We vary the strength of neuronal adaptation, *w*_*A*_, and the intensity of the coherent and incoherent stimuli in each of the relevant directions (0°, 270°, 180°) to test the two hypotheses: stronger adaptation to coherent motion vs. weaker neuronal recruitment in response to coherent stimuli. The first hypothesis predicts that coherent response should be lower than incoherent response only when adaptation is introduced. The second hypothesis predicts that coherent response should be lower than incoherent response regardless of adaptation levels.

### Adaptation is required to replicate experimental order of response curves during stimulus presentation

By varying the adaptation strength parameter (Fig. 3), we observe that increasing adaptation reduces the activity for all conditions during stimulus presentation and inverts the order of the incoherent and non-adapting responses, yielding the experimental order of the curves: coherent < incoherent < non-adapting.

**Figure 3.**
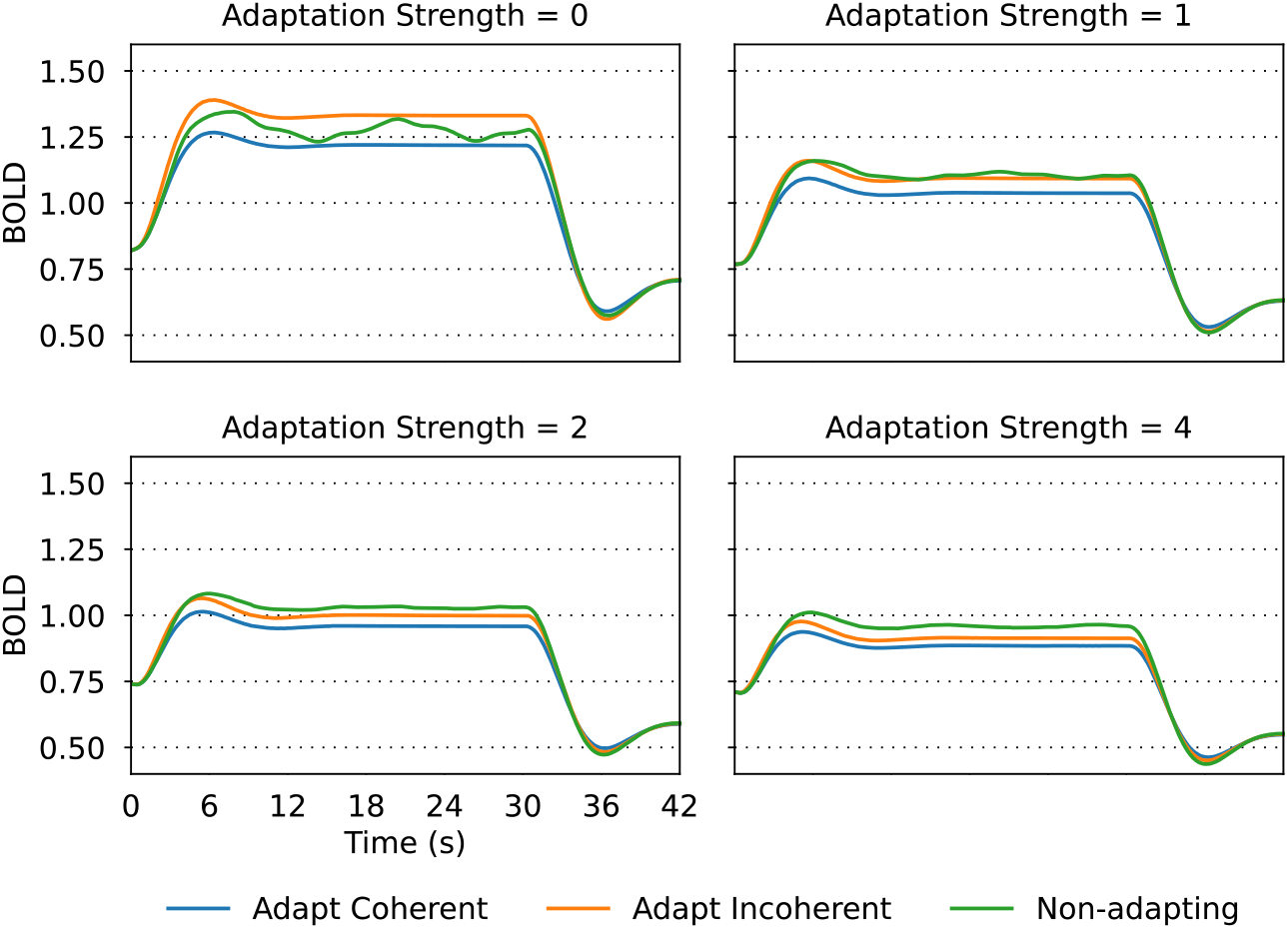
Effect of adaptation strength on the simulated BOLD signal, for each of the experimental conditions (Adapt Coherent, Adapt Incoherent, Non-adapting). Stimulus parameters were *s*_*c*_ = (0,1,0) for coherent motion and *s*_*i*_ = (1/6, 1/3, 1/6) for incoherent motion.

Importantly, the relative order of the coherent and incoherent curves is conserved as adaptation strength increases, suggesting that the lower response to coherent motion stems from the additional mechanism of lower neuronal recruitment, once adaptation is established.

### The model with adaptation also replicates experimental order of response curves during the motion after-affect

After the stimulus is turned off, the order of the curves is reversed relative to stimulus presentation, with the coherent condition eliciting the strongest response (Fig. 4). This agrees with the experimental BOLD signal, as well as the observation of a more vivid MAE following the coherent stimulus, compared to the incoherent condition [18].

**Figure 4.**
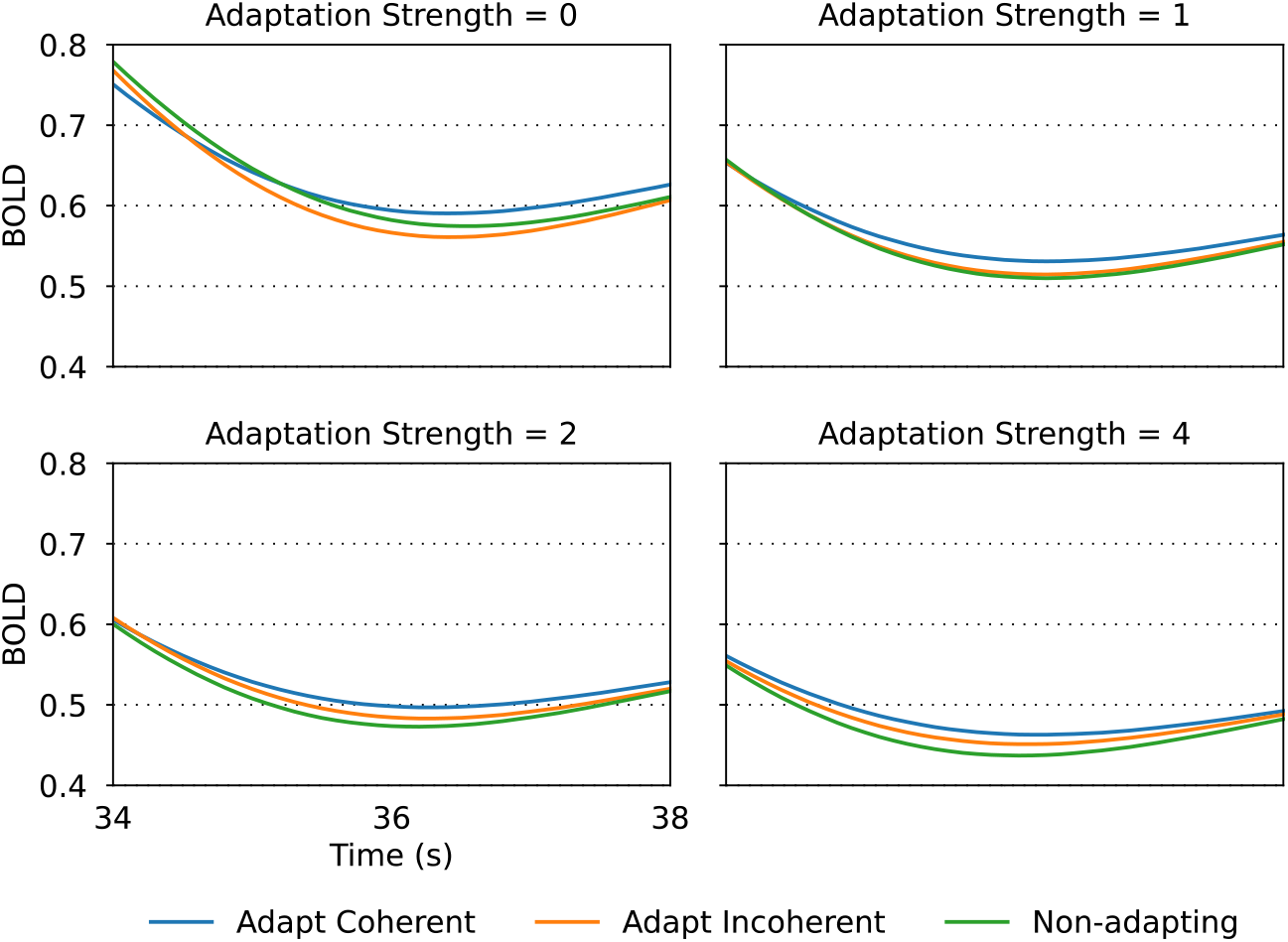
Effect of adaptation strength on the simulated BOLD signal after stimulus presentation, for each of the experimental conditions (Adapt Coherent, Adapt Incoherent, Non-adapting). Stimulus parameters were *s*_*c*_ = (0,1,0) for coherent motion and *s*_*i*_ = (1/6, 1/3, 1/6) for incoherent motion.

### Coherent response is lower than incoherent response across stimulus parametrization and adaptation strength

The definition of the stimulus through the input in each of the three relevant directions (0°, 270°, 180°) is one of the main degrees of freedom of the model. By varying the parametrization of the stimulus (Fig. 5), we can investigate whether it is general that adaptation strength does not affect the relative order of the coherent and incoherent curves, as found in Fig. 3.

**Figure 5.**
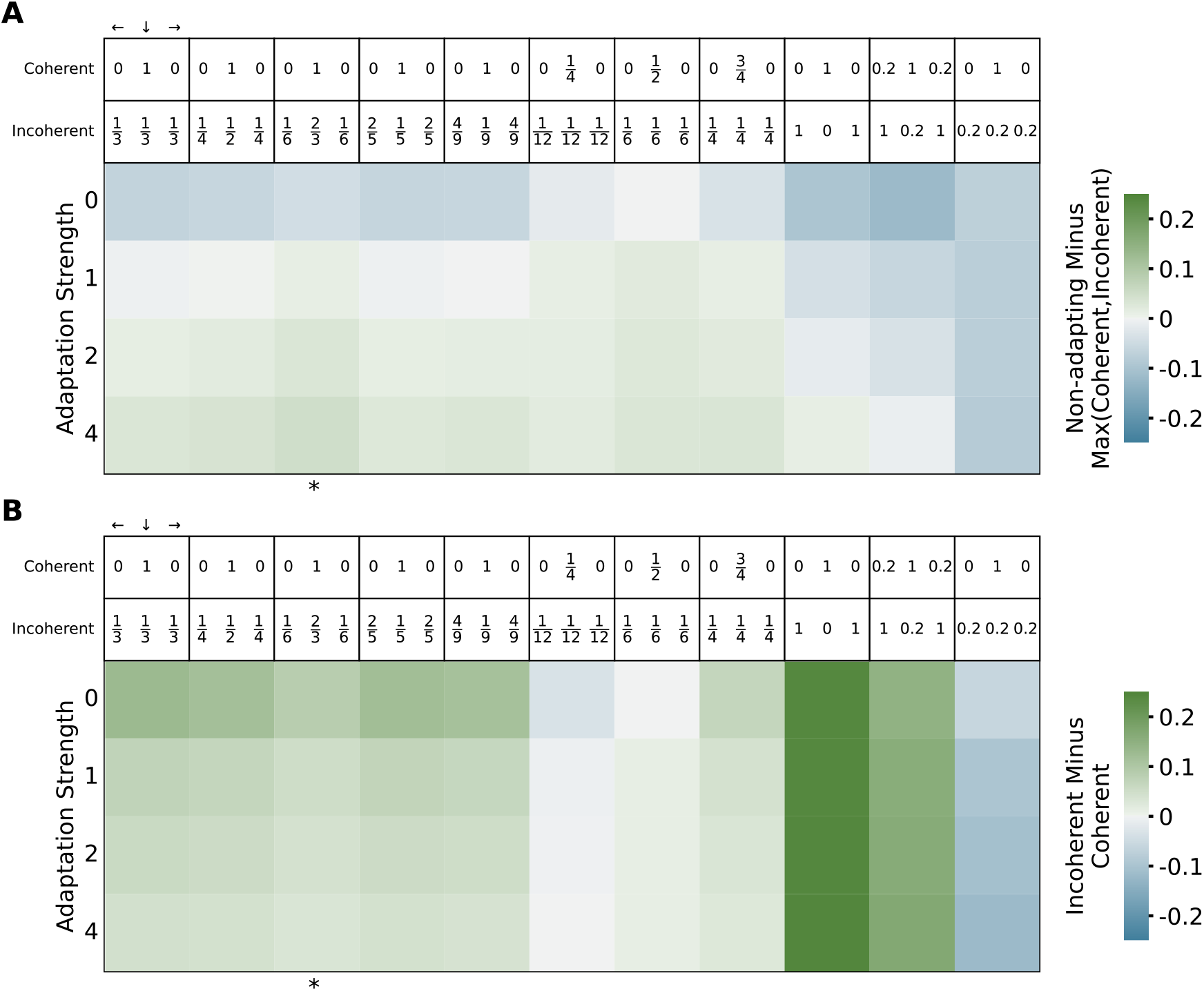
Effect of adaptation strength and different stimulus parametrizations on the relative distance between the curves during stimulus presentation. The metrics were obtained by averaging the point-by-point difference between curves from 6 to 30 seconds. Marked with * is the simulation presented in Figs. 3 and 4. **A**) Positive values (green) denote simulations where the maximum activity is obtained in response to the non-adapting condition. **B)** Positive values (green) correspond to simulations where the response to the coherent condition is weaker than to the incoherent condition. Positive values (green) on both heatmaps represent simulations that replicate the experimental order of the curves.

The first five parametrizations assume that the total stimulus intensity is the same for both coherent and incoherent stimuli and is equal to 1, with variations in the relative weights of the lateral and down directions in the incoherent stimulus. The next three parametrizations assume equal weight of the three directions in the incoherent stimulus but scale down the total intensity in both stimuli. The last three parameter choices do not equalize total intensity between the two stimuli, with the last parametrization chosen so that the coherent stimulus is stronger than the incoherent one.

In general, coherent response is lower than incoherent response regardless of adaptation strength, with three exceptions. When stimulus parameters are chosen with a large motion energy imbalance so that coherent response is stronger than incoherent response in the absence of adaptation (last column: *s*_*c*_ = (0, 1, 0) and *s*_*i*_ = (0.2, 0, 0.2)), introducing adaptation is not enough to invert the order of the coherent and incoherent activations. Similarly, when the total intensity of the stimuli is equal but scaled down to 0.25 (middle column: *s*_*c*_ = (0, 1/4, 0) and *s*_*i*_ = (1/12, 1/12, 1/12)), coherent response is larger than incoherent response regardless of adaptation strength. Finally, when total intensity is scaled down to 0.5 (*s*_*c*_ = (0, 1/2, 0) and *s*_*i*_ = (1/6, 1/6, 1/6)), coherent response is approximately equal to incoherent response in the absence of adaptation and becomes lower with increasing adaptation. This is the only case where adaptation strength alone modulates the response levels in a way that is congruent with the experimental results and with the first hypothesis. However, this constitutes a very specific stimulus parametrization, chosen so that total stimulus intensity is 0.5, which coincides with the inflection point of the neuronal non-linear response function (Eq. 3). It is thus unlikely that this finely-tuned parametrization is representative and biologically accurate. Therefore, for all plausible stimulus parameters, adaptation does not modulate the relative order of the coherent and incoherent response levels.

## Discussion

Visual perception of bistable stimuli has long puzzled scientists and while several mechanisms have been proposed to explain how different interpretations coexist, the specifics have remained elusive. In this work, we focused on a neuroimaging experiment of a bistable plaid stimulus and employed a computational model to study the neural substrate of coherent and incoherent motion perception. Our simulations of motion-sensitive neurons allowed us to reveal the contributions of evoked neural response and adaptation, and how they interact with each other to explain neural responses at a columnar level. Particularly, we tested two alternative hypotheses to explain the lower response to adapting coherent stimuli.

The first hypothesis proposes that coherent stimuli elicit a neural response that matches or surpasses the response to incoherent stimuli in the absence of habituation, but it is adaptation, or repetition suppression, that subsequently diminishes coherent response below incoherent response. This pattern would be evident in simulations with and without neuronal adaptation: in the absence of adaptation, coherent response would be equal to or stronger than incoherent response, whereas with adaptation, coherent response would be lower than incoherent response. Conversely, the second hypothesis states that coherent stimuli elicit a neural response lower than the response to incoherent stimuli, because coherent motion only stimulates neurons tuned to one motion direction, while incoherent motion stimulates at least two neural populations tuned to opposite directions of motion [31]. Simulations would thus show lower coherent response regardless of adaptation levels.

Our results show that only with adaptation can the experimental order of the response curves be replicated, both during stimulus presentation (Fig. 3) and during the motion after-effect (Fig. 4). However, the level of response to coherent motion is generally lower than the response to incoherent motion, irrespective of adaptation strength (Fig. 5), suggesting that differential neuron population recruitment is responsible for the relative order of the coherent and incoherent activations. Hence, the hemodynamic response observed by Sousa et al. [18] arises due to a combination of neuronal adaptation and differences in columnar recruitment.

Our simulations exhibit an important signature of the motion after-effect that occurs after adaptation to a moving stimulus. When the stimulus is turned off, the activity after coherent and incoherent motion is larger than the activity after the non-adapting condition, with adaptation to coherent motion eliciting a stronger after-effect than incoherent motion. It is significant that we replicate the MAE and its neuroimaging signal with our parsimonious model, containing only non-interacting neurons that undergo adaptation.

A noteworthy limitation of our model is its inability to replicate the second half of the BOLD signal recorded by Sousa et al., where the adapting responses increase beyond the non-adapting response [18]. This suggests that other computational motifs may be involved, namely network interactions through inhibition or adapted inhibition [32].

In conclusion, our computational model of motion-sensitive adapting neurons replicates the order of the neural responses to coherent, incoherent, and non-adapting motion observed in an fMRI experiment by Sousa et al. [18], but only when adaptation is present. Furthermore, we tested two competing hypotheses regarding the mechanisms involved in coherent and incoherent motion perception: stronger adaptation to coherent motion vs. stronger neuronal recruitment by incoherent stimuli. By simulating the response of the network to stimuli with various parametrizations and in different adaptation regimes, we determined that the second hypothesis is more likely. Finally, our results also reproduce the neural activity after stimulus exposure, which is congruent with the observed motion after-effect. In sum, this computational work supports the involvement of both adaptation and differential neuronal recruitment as pivotal mechanisms in bistable motion perception.

All data and code used for running simulations, analysis, and plotting are available on Zenodo athttps://doi.org/10.5281/zenodo.10753265.

## References

1. Benda J. Neural adaptation. Curr Biol. 2021;31: R110–R116. doi:10.1016/j.cub.2020.11.054

2. Kohn A. Visual Adaptation: Physiology, Mechanisms, and Functional Benefits. J Neurophysiol. 2007;97:3155–3164. doi:10.1152/jn.00086.2007

3. Wark B, Lundstrom BN, Fairhall A. Sensory adaptation. Curr Opin Neurobiol. 2007;17:423–429. doi:10.1016/j.conb.2007.07.001

4. Webster MA. Visual Adaptation. Annu Rev Vis Sci. 2015;1:547–567. doi:10.1146/annurev-vision-082114-035509

5. Anstis S, Verstraten FAJ, Mather G. The motion aftereffect. Trends Cogn Sci. 1998;2:111–117. doi:10.1016/S1364-6613(98)01142-5

6. Price NSC, Prescott DL . Adaptation to direction statistics modulates perceptual discrimination. J Vis. 2012;12:32–32. doi:10.1167/12.6.32

7. Harvey BM, Braddick OJ . Similar adaptation effects on motion pattern detection and position discrimination tasks: Unusual properties of global and local level motion adaptation. Vision Res. 2011;51:479–488. doi:10.1016/j.visres.2011.01.002

8. Van Wezel RJA, Britten KH . Motion Adaptation in Area MT . J Neurophysiol. 2002;88:3469–3476. doi:10.1152/jn.00276.2002

9. Barlow HB, Hill RM . Evidence for a Physiological Explanation of the Waterfall Phenomenon and Figural After-effects. Nature. 1963;200:1345–1347. doi:10.1038/2001345a0

10. Krekelberg B, Boynton GM, Van Wezel RJA. Adaptation: from single cells to BOLD signals. Trends Neurosci. 2006;29:250–256. doi:10.1016/j.tins.2006.02.008

11. Barron HC, Garvert MM, Behrens TEJ. Repetition suppression: a means to index neural representations using BOLD? Philos Trans R Soc B Biol Sci. 2016;371: 20150355. doi:10.1098/rstb.2015.0355

12. Grill-Spector K, Malach R. fMR-adaptation: a tool for studying the functional properties of human cortical neurons. Acta Psychol (Amst). 2001;107:293–321. doi:10.1016/S0001-6918(01)00019-1

13. Huk AC, Heeger DJ . Pattern-motion responses in human visual cortex. Nat Neurosci. 2002;5:72–75. doi:10.1038/nn774

14. Tootell RBH, Reppas JB, Dale AM, Look RB, Sereno MI, Malach R, et al. Visual motion aftereffect in human cortical area MT revealed by functional magnetic resonance imaging. Nature. 1995;375:139–141. doi:10.1038/375139a0

15. Huk AC, Ress D, Heeger DJ . Neuronal Basis of the Motion Aftereffect Reconsidered. Neuron. 2001;32:161–172. doi:10.1016/S0896-6273(01)00452-4

16. Blake R, Logothetis NK . Visual competition. Nat Rev Neurosci. 2002;3:13–21. doi:10.1038/nrn701

17. Tong F, Meng M, Blake R. Neural bases of binocular rivalry. Trends Cogn Sci. 2006;10:502–511. doi:10.1016/j.tics.2006.09.003

18. Sousa T, Sayal A, Duarte JV, Costa GN, Martins R, Castelo-Branco M. Evidence for distinct levels of neural adaptation to both coherent and incoherently moving visual surfaces in visual area hMT+. NeuroImage. 2018;179:540–547. doi:10.1016/j.neuroimage.2018.06.075

19. Friston KJ, Harrison L, Penny W. Dynamic causal modelling. NeuroImage. 2003;19:1273–1302. doi:10.1016/S1053-8119(03)00202-7

20. Aschner A, Solomon SG, Landy MS, Heeger DJ, Kohn A. Temporal Contingencies Determine Whether Adaptation Strengthens or Weakens Normalization. J Neurosci. 2018;38:10129–10142. doi:10.1523/JNEUROSCI.1131-18.2018

21. Dayan P, Abbott LF . Theoretical neuroscience: computational and mathematical modeling of neural systems. Cambridge, Mass: Massachusetts Institute of Technology Press; 2001.

22. Cravo MI, Bernardes R, Castelo-Branco M. Subtractive adaptation is a more effective and general mechanism in binocular rivalry than divisive adaptation. J Vis. 2023;23: 18. doi:10.1167/jov.23.7.18

23. Gast R, Rose D, Salomon C, Möller HE, Weiskopf N, Knösche TR. PyRates—A Python framework for rate-based neural simulations. Lytton WW, editor. PLOS ONE. 2019;14: e0225900. doi:10.1371/journal.pone.0225900

24. Gast R, Knösche TR, Kennedy A. PyRates—A code-generation tool for modeling dynamical systems in biology and beyond. Marinazzo D, editor. PLOS Comput Biol. 2023;19: e1011761. doi:10.1371/journal.pcbi.1011761

25. Virtanen P, Gommers R, Oliphant TE, Haberland M, Reddy T, Cournapeau D, et al. SciPy 1.0: fundamental algorithms for scientific computing in Python. Nat Methods. 2020;17:261–272. doi:10.1038/s41592-019-0686-2

26. Said CP, Heeger DJ . A Model of Binocular Rivalry and Cross-orientation Suppression. Mamassian P, editor. PLoS Comput Biol. 2013;9: e1002991. doi:10.1371/journal.pcbi.1002991

27. Li H-H, Rankin J, Rinzel J, Carrasco M, Heeger DJ . Attention model of binocular rivalry. Proc Natl Acad Sci. 2017;114. doi:10.1073/pnas.1620475114

28. Shpiro A, Moreno-Bote R, Rubin N, Rinzel J. Balance between noise and adaptation in competition models of perceptual bistability. J Comput Neurosci. 2009;27:37–54. doi:10.1007/s10827-008-0125-3

29. Deco G, Jirsa VK . Ongoing Cortical Activity at Rest: Criticality, Multistability, and Ghost Attractors. J Neurosci. 2012;32:3366–3375. doi:10.1523/JNEUROSCI.2523-11.2012

30. Cakan C, Jajcay N, Obermayer K. neurolib: A Simulation Framework for Whole-Brain Neural Mass Modeling. Cogn Comput. 2021 [cited 31 Jan 2023]. doi:10.1007/s12559-021-09931-9

31. Castelo-Branco M, Formisano E, Backes W, Zanella F, Neuenschwander S, Singer W, et al. Activity patterns in human motion-sensitive areas depend on the interpretation of global motion. Proc Natl Acad Sci. 2002;99:13914–13919. doi:10.1073/pnas.202049999

32. Sousa T, Sayal A, Duarte JV, Costa GN, Castelo-Branco M. A human cortical adaptive mutual inhibition circuit underlying competition for perceptual decision and repetition suppression reversal. NeuroImage. 2024;285: 120488. doi:10.1016/j.neuroimage.2023.120488

